# Interactive effects of Nutrient Enrichment and Water Scarcity on Floral signaling and visitation

**DOI:** 10.64898/2026.11.01.763433

**Authors:** Murugash Manavalan, Dinesh Thakur, Bidhan Panthi, Zuzana Münzbergová

## Abstract

• Floral traits remain an underexplored frontier in global change biology compared to their vegetative counterparts. As anthropogenic change involves simultaneous shifts in resource availability, focusing on isolated stressors may overlook complex biological responses. Characterizing the interactive effects of these stressors is therefore crucial to move beyond additive models and accurately predict the stability of pollination services in a transforming world.

• The individual and interactive effects of nutrient enrichment and water scarcity were investigated on the floral traits and pollinator visitation rates of five grassland species.

• While species identity emerged as the dominant driver of overall trait variation, across- species responses were also present. Among the abiotic drivers, nutrient enrichment emerged as the primary influence, significantly increasing flower number and achromatic contrast while decreasing floral area, chromatic contrast and chroma. Conversely, water scarcity primarily reduced plant height and increased floral chroma and hue. The two factors also often interacted in their effects. The changes in visual traits scaled up to changes in pollinator visitation, but the specific effects were largely species-specific.

• Ultimately, our findings demonstrate that anthropogenic stressors act in divergent ways to reshape floral phenotypes and pollinator visitation, advancing our understanding of plant- pollinator interactions under compounding global pressures.

## 1. Introduction

Angiosperms represent one of the ecologically dominant groups of terrestrial life and today represent ∼90% of all land plants in the globe. A key innovation which underpins this diversification is the greater structural complexity of flowers and the evolution of biotic pollination (Van Der Niet and Johnson 2012, Benton, Wilf, and Sauquet 2022). The crucial mutualism between angiosperms and pollinators is primarily maintained through floral rewards which provide nutrition for pollinators and facilitates plant reproduction (Nicolson *et al*. 2018, Hernández-Hernández and Wiens 2020). These vital interactions which co-evolved over millennia now face growing threats due to strong fast-acting anthropogenic pressures (Vanbergen and Initiative 2013). These pressures render the diversity and abundance of flowering plants under constant threat with cascading consequences for pollinators (Martén Rodríguez *et al*. 2025). Given that flowers are the primary medium of interaction between plants and pollinators, understanding how flowers respond to stressors becomes essential for assessing broader implications for plant-pollinator interactions, subsequent ecosystem processes and services dependent on plant-pollinator interactions.

Two major anthropogenic stressors which are expected to have major impacts on plant reproduction and pollinators are nutrient enrichment which occurs through agricultural runoff and atmospheric deposition, and water scarcity which is driven by extensive land-use and climate change (Foley *et al*. 2005, Isbell *et al*. 2013, Crausbay *et al*. 2017). These stressors tend to elicit contrasting responses on flowers with nutrient enrichment found to largely exert positive effects on floral traits (Cardoza, Harris, and Grozinger 2012, Atasay *et al*. 2013, Vaudo *et al*. 2022) while water scarcity is predominantly associated with negative outcomes (Snider and Oosterhuis 2011, Rering *et al*. 2020, Kuppler and Kotowska 2021). Different environmental stressors however rarely act in isolation, and their combined effects often produce complex, non-additive outcomes that single-factor studies fail to capture (Atkinson and Urwin 2012, Münzbergová *et al*. 2017, Rongstock *et al*. 2026). Studies exploring interactive effect of different environmental factors on plant flowers and plant-pollinator interactions are however rare. In an exceptional study, Akter and Klečka 2022 demonstrated that water and nitrogen availability indeed interact with each other to affect nectar volumes in plants. How these factors interact to affect other flower traits relevant to plant pollinator interactions remain to be explored. Importantly, flower color is important to aid in pollinator attraction but also play key roles in plant’s physiological response to abiotic stress (Koski and Ashman 2015, Kooi van der *et al*. 2019). Responses of flower color to environmental changes are however largely unexplored (Nelson, Borer, and Seabloom 2025).

There is growing evidence that anthropogenic activities have triggered widespread declines in wild pollinator populations (Vanbergen and Initiative 2013, Dicks *et al*. 2021). However, experimental studies investigating the downstream consequences of stress-induced floral traits shifts on pollinator visitation remain critically limited and highly variable in their outcomes. While some studies have investigated the effect of drought on plant-pollinator interactions through changes in floral traits (reviewed in Kuppler and Kotowska 2021), the effect of interacting abiotic factors still remains relatively unexplored given the methodological complexity behind it (but see Glenny, Runyon, and Burkle 2018). By explicitly bridging the gap between interacting abiotic stressors, altered floral traits and pollinator behaviour, we could better understand how climate change and habitat modification can affect pollination services and ecosystem function.

Most of the above cited studies exploring the effects of different environmental stressors on plant flowers and plant-pollinator interactions are restricted to single plant species. While metanalyses such as (Kuppler and Kotowska 2021) are generally useful to summarize effects across studies, they are unlikely to find any strong generalities due to different levels of the different factors studied and the effects often being context dependent. Studies exploring the effects of the same treatments across different species are thus needed to allow for any generalizations but are largely missing.

We aim to fill the above gaps by exploring the effects of nitrogen and water availability and their interaction on plant flower traits including the largely unstudied flower color and explicitly testing how these trait changes translate to pollinator visitations. By performing this study across 5 different co-occurring species from different families under identical, standardized conditions, we also aim to assess how general these patterns are.

Specifically, we ask the following questions:

**Q1.** What are the effects of nutrient enrichment, water scarcity, and their interaction on floral visual traits across different species? **Q2.** How do nutrient enrichment, water scarcity, and their interaction influence pollinator visitation? **Q3.** How do these phenotypic changes and their role in pollinator attraction vary among species?

We hypothesize that **H1.** Across species, nutrient enrichment will exert overall positive effects on floral visual traits, whereas water scarcity will have negative effects. Under combined treatment, nutrient enrichment will partially mitigate the negative impacts of water deficit by improving drought tolerance. **H2.** Pollinator visitation will be driven by shifts in floral traits caused by the abiotic treatments. Specifically, nutrient enrichment will increase flower visitation by enhancing visual display traits, while water scarcity will decrease attraction. **H3.** Species- specific responses will mirror the ecological strategies of their native environments, reflecting distinct trade-offs between stress tolerance and resource acquisition strategies. Consequently, the specific traits dictating pollinator attraction will vary by plant species, and the phenotypic plasticity of species-specific floral traits under stress will ultimately determine the magnitude of change in pollinator visitation rates.

## 2. Materials and Methods

### 2.1. Plant species and experimental design

Five perennial entomophilous species specific to dry and mesic grasslands of Europe were selected for this study: *Arabidopsis arenosa* (L.) Lawalrée (Brassicaceae), *Dianthus deltoides* L. (Caryophyllaceae), *Leucanthemum vulgare* subsp. *ircutianum* (Turcz. ex DC.) Tzvelev (hereafter referred to as *L. ircutianum*) (Asteraceae), *Potentilla argentea* L. (Rosaceae) and *Veronica chamaedrys* L. (Plantaginaceae). These five species were chosen as they were the only ones within a larger multi-species experiment that successfully reached full anthesis and flowered across all experimental treatments. Seeds were acquired from a commercial supplier (Planta Naturalis, Markvartice, Czech Republic). Soil to be used was obtained from a nutrient-poor ruderal habitat close to the experimental garden. In summer of 2023, ∼100 seeds/species were sown in a soil mixture containing 1:1 (v/v) soil and sterile sand. Once at least 50 seedlings germinated, the individual seedlings were transferred to multi-pots with the same mixture of soil and sand where they were allowed to grow further.

After 2–3 weeks, 36 individuals of each species were transplanted into 2l pots and assigned to four treatments (each n = 9 plants): natural soil with regular water supply (Poor-Wet, PW), fertilized soil with regular water supply (Rich-Wet, RW), natural soil with restricted water supply (Poor-Dry, PD), and fertilized soil with restricted water supply (Rich-Dry, RD). For the fertilized soil treatments, 6g of slow-release fertilizer pellets (N:P:K 19:9:10 + 2 MgO + Trace elements, Osmocote® Pro, Everris International, Geldermalsen, the Netherlands) was mixed in with the soil. The amount of fertilizer used is in relation to other studies on wild plants (Dawson, Fischer, and Van Kleunen 2012, Klinerová and Dostál 2020). For the regular water supply treatments, plants were kept in open sheds with unrestricted access to rain while also being watered regularly. For the restricted water supply treatment, the plants were categorized into 3 groups based on their Ellenberg Indicator Moisture value (Chytrý *et al*. 2018): *P. argentea* in group “2”, *A. arenosa* and *D. deltoides* in group “4” and finally *L. ircutianum* and *V. chamaedrys* in group “5”. This type of categorization is based on a reliable indicator system (Schaffers and Sýkora 2000) which takes into account the prevailing moisture conditions in which the plants naturally occur and thus allowed us to impose water restriction on all plants considering their differing water requirements. The water restriction treatment was such that, when one individual of a certain moisture group began to show signs of wilting, then all of the plants in that group irrespective of species were watered. To monitor the moisture values of the treatments, TMS dataloggers were used where each moisture group in the restricted water supply treatment was assigned one datalogger while two dataloggers were assigned for all the plants in the regular water supply treatment (TOMST s.r.o, Czech Republic) (Wild *et al*. 2019). Plants were grown under treatment conditions through the summer and autumn of 2023. Treatments were suspended during winter of 2023-2024 to minimize overwinter mortality and were resumed in spring 2024.

### 2.2. Morphological and Reflectance measurements

Beginning in spring 2024, one month following the resumption of treatment conditions, floral morphological traits related to pollinator attraction, specifically floral height, flower number and floral area were measured for all plants using standardized protocols (Pérez-Harguindeguy *et al*. 2013, Michelot Antalik *et al*. 2025). It should be noted that in *L. ircutianum*, a flower refers to one inflorescence composed of both ligulate and tubular florets. Using a standard ruler, floral height was taken as the shortest distance between the top-most flower of an individual and the ground (in cm). Flower number was taken as the number of antithetical open flowers (or flowerheads in *L. ircutianum*) and buds ready to bloom in an individual. Flower number for a species was recorded when atleast two out of the four treatments were in their peak flowering stage. For the floral area, calipers were used to measure the diameter (d, to the nearest mm) of one flower per individual following which the area was calculated according to the equation A = π(d/2)^2^.

Floral reflectance data was acquired from one flower per individual (ligulate florets from *L. ircutianum*) using a AVASpec-2048 fiber-optic spectrometer (Avantes Starline, Netherlands) equipped with a xenon light source (Avalight-XE-HP, Avantes) and a black metal probe holder (6.5mm diameter opening 45°)(RPH-1, Avantes). All measurements were done in a black chamber. The spectrometer was calibrated both with white standard (WS-2, Avantes) and total absence of light. Petals were placed on black insulation tape, and their reflectance curves were measured from 200-1100nm wavelengths at 0.6nm intervals with an integration time of 5s per measurement. The distance between the optic fiber and the petals was kept consistent across all measurements for each species with three measurements taken per individual (Michelot Antalik *et al*. 2025).

Analysis of the reflectance measurements were done following the methodology of Del Valle *et al*., 2025. Wavelengths between 300 and 700m were considered (Endler and Mielke 2005, Del Valle *et al*. 2025). After importing the reflectance spectra into R using the ‘pavo’ package (v. 2.9; Maia *et al*., 2019), the *procspec* function was used to smooth the reflectance curves with a smoothness parameter of 0.2 and remove negative values by setting minimum values to zero and scaling other values accordingly (Del Valle *et al*. 2025). Animal vision models of both hymenopterans (bees) and dipterans (flies) were used to represent how a colour stimulus is perceived by the two dominant groups of pollinators. **Chromatic contrast** was calculated as the Euclidean distance between the colour loci of the flower and the achromatic centre measured in Euclidean units (EU). For bees, **achromatic contrast** was calculated as the difference between the excitation value of the green photoreceptor and 0.5, which represents adaptation to the background (Del Valle *et al*. 2025). Other spectral traits including **total brightness, chroma** and **hue** were also calculated using *summary.rspec* function in the ‘pavo’ package (v. 2.9; Maia *et al*., 2019). Total Brightness is taken as the sum of reflectance (%) from 300-700nm. Chroma is calculated as (Rmax – Rmin)/Rmean where R is reflectance (nm). Hue corresponds to the wavelength of maximum reflectance (Michelot Antalik *et al*. 2025).

### 2.5. Behavioral assays

To assess the effects of soil nutrient and water treatments on flower visitation, a factorial flower visitor preference experiment was conducted for each plant species. Pollination experiments were only carried out once all four treatments of a species achieved peak flowering and care was taken to minimize interval between measurement of floral traits and behavioral assays to ensure parity between the two sets of variables. Here, peak flowering is defined as plant individuals having more flowers in anthesis stage than in budding stage. Each of the four treatments was paired with every other treatment, resulting in six unique treatment combinations (PD × RD, PD × PW, PD × RW, RD × PW, RD × RW, and PW × RW). Each treatment combination (henceforth referred to as block) was observed for 20 minutes, yielding a total observation time of 2 hours per species.

Experiments were conducted at a site separate from the plant cultivation area to ensure unrestricted access by natural flower visitor communities and to minimize potential carryover effects from the growing environment. For each experimental block, eight flowering individuals from each treatment were selected and arranged alongside individuals from the paired treatment, with plant positions randomized to avoid spatial bias (Illustrated in Fig. 1). Pots were placed 20 cm apart to allow clear separation between individuals while approximating natural foraging conditions. Environmental conditions were monitored throughout the trials. Wind speed, a key factor influencing pollinator activity, was measured using an anemometer, and observations were conducted only when wind speeds did not exceed 3.5 m s ¹. Temperature, another key metric was recorded by means of Weather Apps with care being taken to only undertake observations when ambient temperatures were between 15 - 25 °C. These considerations alongside ensuring that experiments only took place under clear sky conditions between 09:00 and 17:00 h ensured favorable and consistent conditions for pollinator activity.

**Fig. 1.**
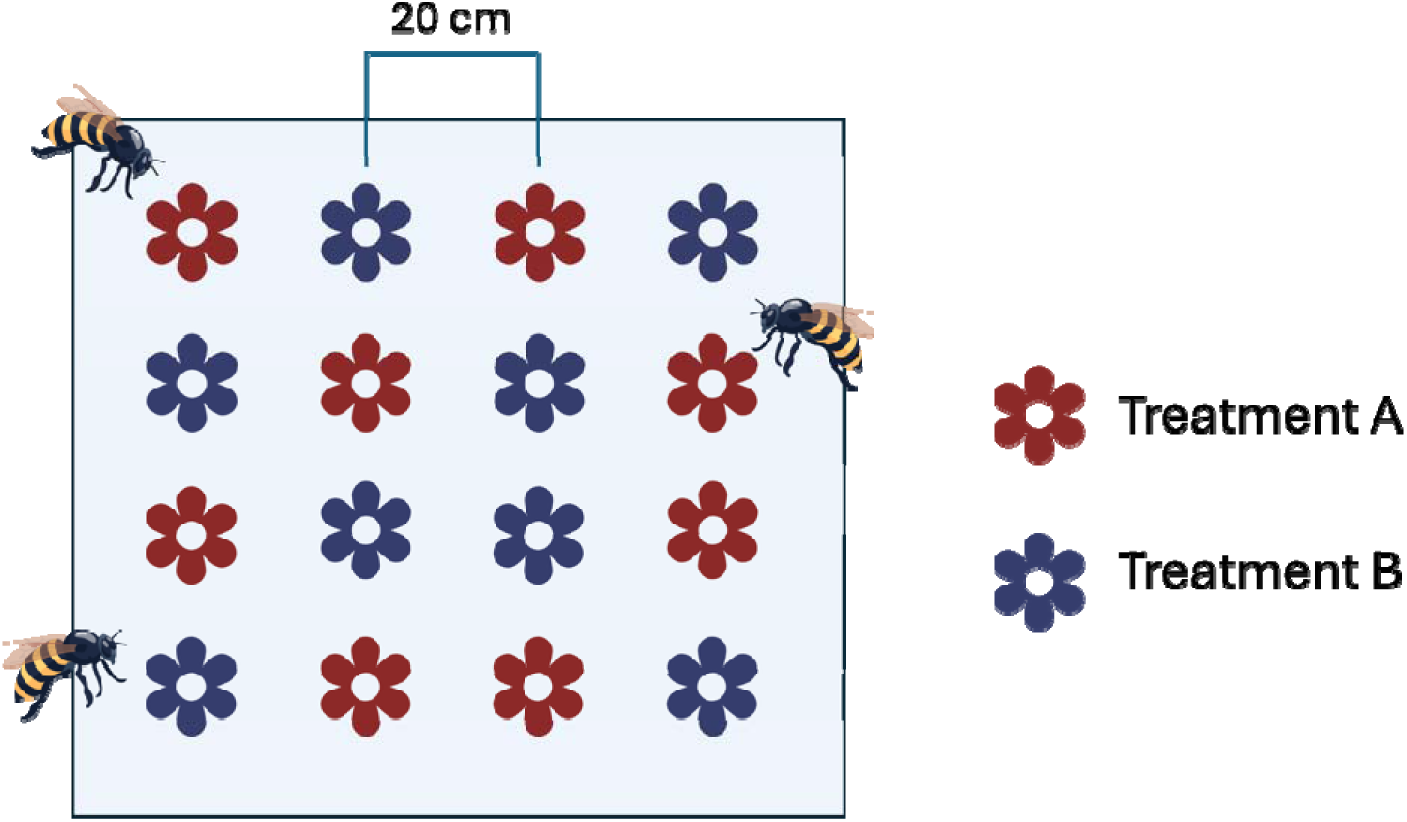
Experimental setup for pollinator visitation. Each experimental block consists of two treatments with equal number of individuals from both treatments randomly arranged in a field with adequate presence of pollinators. Icons from acquired from Flaticon.

Although the reproductive efficacy of floral visitors was not directly evaluated in this study, all visitors are hereafter referred to as ‘pollinators’. During each block observation period, the pollinator visitation rate was recorded as the total number of visits received by a plant individual. A visit was defined as an insect landing on a flower of a plant individual; consecutive visits to multiple flowers on the same individual were recorded as a single visit. However, if a pollinator visited a different individual before returning to the initial plant, this was counted as a distinct visit. Pollinators were categorized into four functional groups: honeybees, bumblebees, solitary bees, and hoverflies.

### 2.6. Statistical analyses

All statistical analyses were carried out using R programming language (R Core Team 2025). Plots were created using functions from the ggplot2 package (Wickham 2016). AI assistance wa used to generate R code for certain analytical approaches.

#### 2.6.1. Effects of treatments on plant traits

For assessment of impact of factors on individual plant traits across all species, a Shapiro Wilk test was first done to evaluate the normality of trait values. The normality of the original trait value was compared with its logarithmic and square-root transformations based on the P-values obtained. The form of trait-value with the highest P-value was selected for inclusion in linear models. Plant Height, Chromatic Contrast (for bees and flies), Achromatic Contrast, Total Brightness and Hue were kept in their original form; Floral Area, and Chroma were log- transformed while Flower Number was square-root transformed. Linear models were used to analyze the effects nutrient enrichment, water scarcity and their interaction on floral traits. These models were used in two levels: 1. Across all species, to assess general patterns in trait responses to nutrient enrichment, water limitation, their interaction and interaction of all these with species; and 2. Separately per species in cases where species x treatment interactions are detected. Analysis of variance (ANOVA) was used to assess statistical significance on each floral trait. Where ANOVA indicated a significant effect of interaction between nutrient enrichment and water scarcity in a particular species, Tukey’s Honestly Significant Difference (HSD) post hoc test was used to identify which specific treatment combinations differed significantly from one another. To understand relationship between individual traits, correlation heatmaps were created with the corrplot package (v.0.95; Wei and Simko, 2024).

#### 2.6.2. Effect of abiotic factors on flower visitation

While we recorded the functional groups per block in order to assess the effect of abiotic factors on pollinator diversity, lack of visits of different groups across multiple blocks made this an unviable option. The effects of abiotic factors on flower visitation were analyzed using generalized linear mixed-effects models (GLMMs) with a negative binomial error distribution. The negative binomial distribution was selected over a Poisson distribution to account for overdispersion in pollinator visitation counts, where the variance exceeded the mean. Nutrient treatment, water treatment, species identity, and their interactions were included as fixed effects, together with environmental covariates (cloud cover, wind speed, and temperature). Experimental block was included as a random intercept to account for using the same individuals arranged in different blocks for pollinator observations.

#### 2.6.3. Structural Equation Modelling (SEM)

To disentangle the effects of abiotic factors on flower visitor responses while accounting for intercorrelations among floral visual traits, piecewise structural equation modelling was employed for both across species and also independently for each of the five study species using the ’piecewiseSEM’ package (v. 2.3.1; Lefcheck, 2016). All traits were standardized (mean = 0, SD = 1) prior to analysis. The model structure followed a three-level hierarchy: nutrient and water treatments and their interaction were specified as exogenous variables; floral visual traits were modelled as mediatory endogenous variables responding to these treatments and finally flower visitation was modelled as the terminal endogenous variable.

Trait sub-models were fitted to characterize the effects of abiotic factors and their interaction on each floral visual trait, with trait values transformed as previously described. A visitation response model was then fitted with abiotic factors, their interaction, and floral visual traits as predictors of flower visitation. Trait sub-models and visitor response models were fitted on two separate datasets reflecting the structure of the experimental design. Trait values were recorded once per individual plant and analyzed at the plant level. Flower visitation, however, was recorded within a blocked design in which each plant appeared across three separate blocks, each contrasting it against plants from a different treatment combination. In both the trait sub-models and the visitation response model, only terms at least marginally significant at p < 0.1 were retained. Models with no-significant terms were excluded from the final SEM. It should be noted that the SEM representing all species was done using residuals of trait and pollinator visitation values derived from linear models accounting for species identity (Trait/Pollinator visitation ∼ Species). The final SEM for each group (across species and individual species) was assembled from the trait sub-models, the visitation response model and a set of correlated residual terms connecting pairs of traits expected to co-vary due to shared developmental and spectral constraints. As the behavioral assays were conducted at a separate site away from the experimental garden, any remaining direct effects of abiotic factors on flower visitation are likely attributable to unmeasured plant traits not included in this study. It should be noted that the direction of the path coefficient for the interaction between nutrient enrichment and water scarcity must be interpreted within its conditional context: a positive (+) coefficient can result from trait shifts induced by either the simultaneous presence or absence of both factors, whereas a negative (-) coefficient indicates effects driven by the presence of a single abiotic factor. Therefore, SEM outputs for interaction pathways were evaluated alongside violin plots from linear models to clarify how nutrient enrichment and water scarcity interactively affected floral traits. Model fit was assessed using Fischer’s C statistic where a non-significant Fischer’s C (p > 0.05) was taken as indicating adequate fit. Where d-separation tests identified significant conditional dependencies (p < 0.05), missing paths were added and the models were refitted until no additional conditional dependencies were detected.

## 3. Results

### 3.1. Trait Responses to Treatments

#### 3.1.1. Overall patterns

Utilizing the combination of linear models (Fig. S1), variance partitioning of predictors (Fig. 2) and SEM (Fig. 3a), it was found that across all species, nutrient enrichment emerged as the primary abiotic predictor of plant trait variation. Nutrient enrichment was associated with significant increases in flower number and achromatic contrast while leading to decreases in floral area, chromatic contrast (as perceived by flies) and chroma (Fig. S1a-e). Although water scarcity affected fewer traits than nutrient enrichment, it still induced significant variation, specifically reducing plant height while increasing chroma and hue (Fig. S1f-h). Among all floral traits investigated, chroma was the sole trait to show a significant response to the interaction between nutrient enrichment and water scarcity. Under nutrient-enriched conditions, chroma was found to be greater in flowers of water-deprived plants relative to well-watered plants (Fig. S1i).

**Fig. 2.**
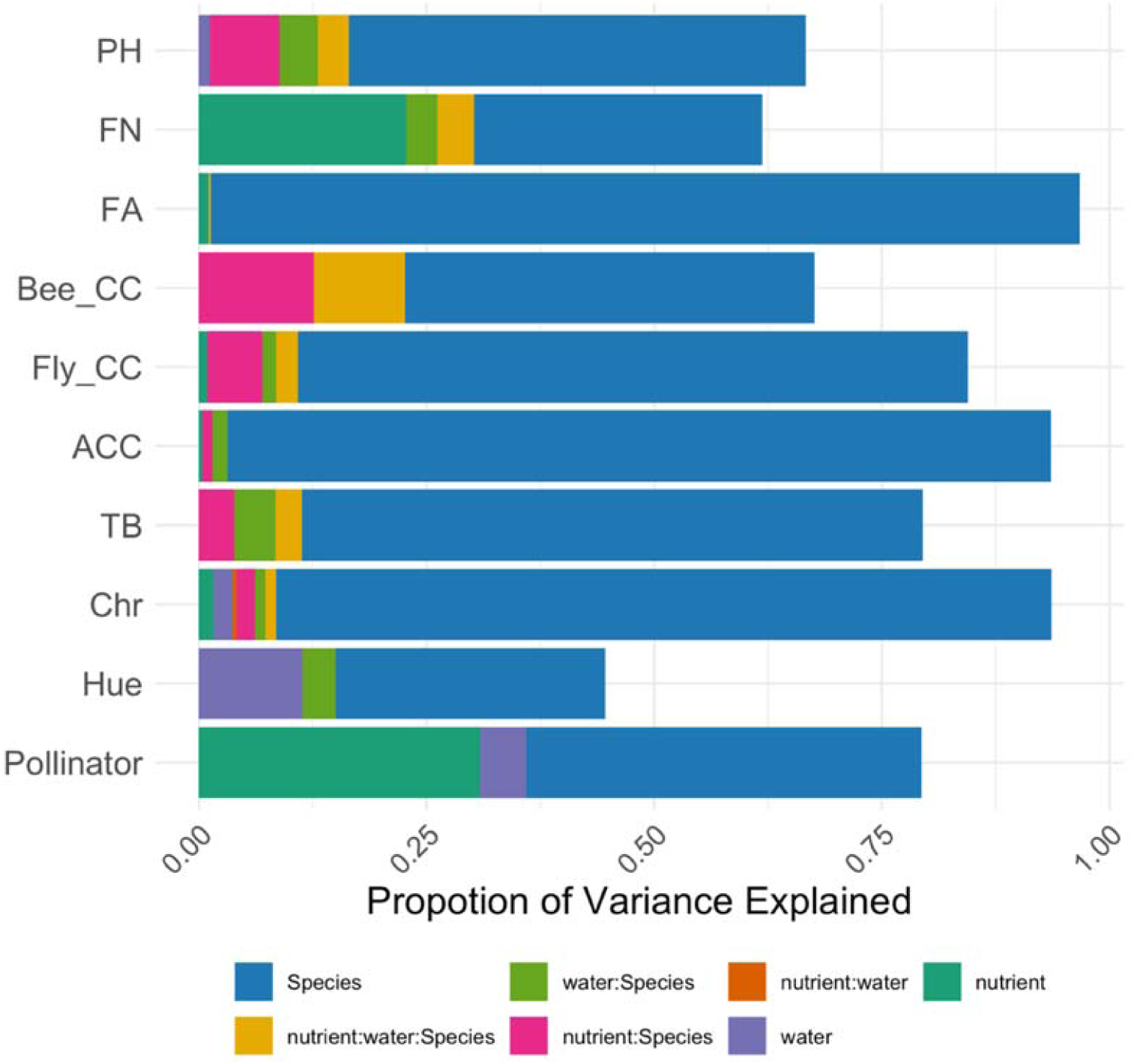
Variance partitioning plot for all species showing magnitude of variation explained by the different abiotic factors. PH, Plant Height; FN, Flower Number; FA, Floral Area; Bee_CC, Chromatic Contrast for Bees; Fly_CC, Chromatic Contrast for Flies; TB, Total Brightness; Chr, Chroma; Pollinator, Pollinator Visitation Rate. Variance explained only by significant variables (p < 0.05) is shown.

**Fig. 3.**
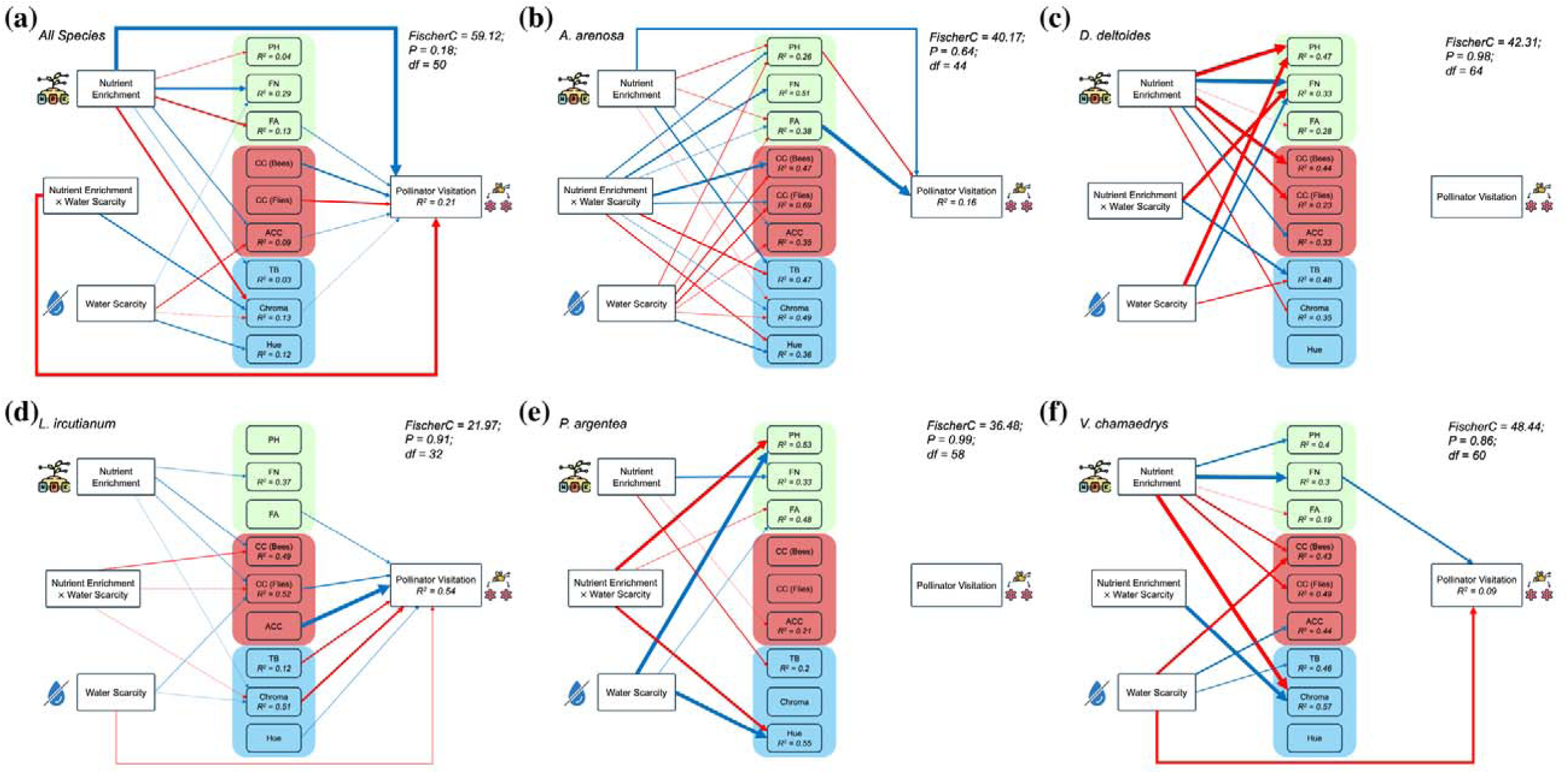
Graphical representation of statistically significant paths retained in final species-specific SEM models. Standardized path coefficients and their level of significance are shown next to the arrows. Blue and red lines represent positive and negative effects respectively. Arrow thickness is proportional to the magnitude of the standardized path coefficient, scaled linearly between the minimum and maximum absolute coefficient values within each plot; thicker arrows therefore indicate stronger effects relative to other paths in the same model. Traits have been grouped into three groups with different colours: morphological traits in green, color contrast traits in red and spectral traits in red. Icons from acquired from flaticon. (a) All Species, (b) *Arabidopsis arenosa*, (c) *Dianthus deltoides,* (d) *Leucanthemum vulgare* subsp. *ircutianum*, (e) *Potentilla argentea*, (f) *Veronica chamaedrys*.

While these general patterns provide valuable insights across species, variance partitioning of predictors (Fig. 2) revealed that species identity was the most influential driver of floral trait variation. Furthermore, the interaction between species identity and abiotic factors further explained substantial variation across nearly all traits, except for hue. This strong context- dependency underscores the need for evaluating and discussing these trait responses at the species-specific level.

#### 3.1.2. Species-specific responses

The below description of the species-specific responses is based primarily on structural equation models (Fig. 3). The figures of the specific patterns as seen from single trait tests and associated violin plots can be found in Supporting Information Fig. S1 – S7 with effects of interaction being detailed in Table S2.

##### a. Arabidopsis arenosa

In *A. arenosa*, nutrient enrichment reduced chroma while increasing total brightness and achromatic contrast. Water scarcity reduced chromatic contrast (in bees and flies), achromatic contrast and chroma, while increasing hue. The interaction between nutrient enrichment and water scarcity produced antagonistic effects on traits relative to the individual treatments, increasing chromatic contrast for both visual systems while decreasing total brightness and hue. Notably, the interaction also produced a significant increase in flower number, a trait that remained unaffected by either factor in isolation (Fig. 3b).

##### b. Dianthus deltoides

In *D. deltoides*, nutrient enrichment reduced plant height, floral area, chromatic contrast for both visual systems, and chroma, while increasing flower number and achromatic contrast. Water scarcity had a more limited effect, reducing plant height and total brightness while unexpectedly increasing flower number. The interaction between the two factors produced antagonistic effects relative to the individual treatments, decreasing flower number and increasing total brightness (Fig. 3c).

##### c. Leucanthemum ircutianum

In *L. ircutianum*, there were very limited trait responses independent of pollinator visitation mediating pathways where nutrient enrichment increased flower number and chromatic contrast for the bee visual system, with the latter decreasing under the combined treatment (Fig. 3d).

##### d. Potentilla argentea

In *P. argentea*, nutrient enrichment decreased achromatic contrast and total brightness while increasing flower number. Water scarcity increased plant height and hue while decreasing floral area. The interaction produced antagonistic effects relative to water scarcity in isolation, decreasing plant height and hue while increasing floral area (Fig. 3e).

##### e. Veronica chamaedrys

In *V. chamaedrys*, nutrient enrichment increased plant height while decreasing floral area, chromatic contrast for both bee and fly visual systems, and chroma. Water scarcity decreased chromatic contrast for the bee visual system while increasing achromatic contrast and total brightness. The interaction produced an antagonistic effect relative to nutrient enrichment in isolation, increasing chroma despite its reduction under nutrient enrichment alone (Fig. 3f).

### 3.2. Effects of Abiotic factors on Pollinator Visitation

#### 3.2.1. Overall patterns

Utilizing SEM, it was found that across species, both abiotic factors and their interaction affected pollinator visitation both directly and indirectly through plant traits (Fig. 3a). Nutrient enrichment while having a strong direct positive effect on pollinator visitation also influenced pollinator visitation through floral area, achromatic contrast and chroma. Although water scarcity was found to not have direct effect on pollination visitation, it indirectly affected pollinator visitation through variation in achromatic contrast and chroma. The interaction between nutrient enrichment and water scarcity was found to influence pollinator visitation through variation in chroma alongside having a direct negative effect. Similar to trait responses to abiotic factors, variance partitioning (Fig. 2) revealed that species identity played a key role in dictating variation in pollinator visitation.

#### 3.2.2. Species-specific variation in pollinator visitation

Between individual species, the effects of abiotic factors on pollinator visitation was only apparent in three out of five species with two species, *D. deltoides* and *P. argentea* lacking any significant link connecting abiotic factors or plant traits to pollinator visitation (Fig. 3c and e). Such links in the remaining three species are elaborated below:

##### a. Arabidopsis arenosa

In *A. arenosa*, both abiotic factors and their interaction affected flower visitation through plant height and floral area where nutrient enrichment and water scarcity in isolation negatively affected both traits while their interaction surprisingly had an antagonistic effect increasing both trait values. Plant height had a negative effect on flower visitation while floral area had a positive effect. Beyond these trait-mediated pathways, nutrient enrichment also exerted a direct positive effect on flower visitation, independent of the measured traits (Fig. 3b).

##### b. Leucanthemum ircutianum

In *L. ircutianum*, pollinator visitation was primarily driven by traits that showed no response to the abiotic treatments. Floral area, achromatic contrast, and hue each had a positive effect on flower visitation, while total brightness had a negative effect. None of these traits were influenced by nutrient enrichment, water scarcity, or their interaction, indicating that the main drivers of visitation in this species operated independently of the experimental treatments (Fig. 3d).

Nevertheless, abiotic factors did influence flower visitation through a separate set of trait- mediated pathways. Both nutrient enrichment and water scarcity, when applied in isolation, positively affected chromatic contrast for the fly visual system and chroma, which in turn had positive and negative effects on flower visitation, respectively. As observed in *A. arenosa*, the interaction of nutrient enrichment and water scarcity produced antagonistic effects relative to the individual treatments, decreasing both chromatic contrast for the fly visual system and chroma. Beyond these trait-mediated pathways, water scarcity also exerted a direct negative effect on flower visitation, independent of the measured traits.

##### c. Veronica chamaedrys

In *V. chamaedrys*, pollinator visitation was driven by two pathways. Nutrient enrichment influenced visitation indirectly through flower number, which it increased, and flower number in turn had a positive effect on visitation. Water scarcity, by contrast, exerted a direct negative effect on flower visitation, independent of the measured traits (Fig. 3f).

## 4. Discussion

This study aimed to characterize how the independent and interactive effects of nutrient enrichment and water scarcity influence floral traits, and how these trait-level responses subsequently shape pollinator visitation. The results indicate that while nutrients, water availability as well as their interaction indeed significantly affect the plant traits, the direction of the effects differed between species making the results hard to generalize. Rather than a single overarching trend, the interaction between both factors operated through two pathways: while nutrient enrichment exhibited a mitigatory effect on negative effects of water scarcity for specific traits in one species (*A.* arenosa), in several instances the influence of nutrient enrichment was dependent on soil moisture availability. Finally, our results supported the idea that some flower traits can be used to explain the plant pollinator interactions, but these effects have also been largely species specific.

### 4.1. Floral trait responses to abiotic stressors

#### 4.1.1. Across-species patterns

Among the floral traits analyzed in this study, flower number was the most responsive to abiotic factors. The consistent stimulation of flower production by nutrient enrichment across most species supports the idea that increased nutrient availability relaxes resource limitation on reproduction, allowing plants to allocate more resources to flowering. Similar responses have been reported across a wide range of species (e.g., Burkle and Irwin 2009, 2010, Leach *et al*. 2023, Wu *et al*. 2023). Nutrient enrichment also led to decrease in chroma and/or chromatic contrast in most of our SEM models. Chroma is closely associated with anthocyanin pigment concentration, with greater anthocyanin content corresponding with higher chroma (Hopkins and Rausher 2012). While abiotic stressors such as nutrient deficiency and drought typically induce anthocyanin production as an adaptive response to oxidative stress (Dabravolski and Isayenkov 2023, Jezek *et al*. 2023), excessive nitrogen fertilization has been found to impair anthocyanin biosynthesis in fruits (Delgado, González, and Martín 2006). The same could have happened in the flowers in our study. In addition to these quite general patterns, most other responses have been species specific and are thus described below.

#### 4.1.2. Individual species patterns

In comparison to the widespread effect of nutrient enrichment on flower number across species, water scarcity was found to have a significant effect on flower number in only one species, *D. deltoides*, where it surprisingly caused an increase. While water scarcity typically reduces floral abundance globally (reviewed in Kuppler and Kotowska 2021, Brunet *et al*. 2025), high heterogeneity exists among species, with some plants producing more flowers under drought (e.g., Campbell and Wendlandt 2013, Walter 2020) which can be viewed as a potential drought- escape strategy in perennial species.

Aside from flower number, water scarcity affected coloration of *A. arenosa*, where it induced a significant decrease in chroma, achromatic contrast, and chromatic contrast (as perceived by bees and flies). This plastic decline in color intensity stands in stark contrast to macroecological trends. For instance, a large-scale study spanning 339 species found that flower color tends to be more chromatic in regions characterized by high solar radiation and low precipitation (Dalrymple *et al*. 2020). However, our observed increase in hue may reconcile these differences. This hue shift reflects a transition towards redder wavelengths, which is a known physiological signature of anthocyanin accumulation under stress (Davies *et al*. 2022).

*A. arenosa* also showed reduced height and floral area under nutrient enrichment. While this seems counter-intuitive, this can be better interpreted when considering its ecological nutrient threshold (reflected in Ellenberg indicator values; Chytrý *et al*. 2018). For *A. arenosa*, a plant adapted to nutrient poor environments (Ellenberg indicator nutrient value = 3), fertilizer application in the nutrient enrichment treatment may have reached toxic thresholds. Nutrient toxicity has been previously known to impair growth and physiology in certain plant species (e.g., Farhan *et al*. 2024, Sarwar *et al*. 2025).

Strikingly, for plant height, chroma and chromatic contrast (as perceived by both bees and flies), nutrient enrichment exhibited a mitigatory effect on the negative impacts of water scarcity. These mitigatory responses are likely driven by nutrient-mediated enhancements in photosynthetic rates, water-use efficiency, and cell membrane integrity under drought conditions (Wu ZZ *et al*. 2018). Conversely, we also observed instances where the physiological influence of nutrient enrichment was dependent on soil moisture conditions. This was evident for total brightness in *A. arenosa* and *D. deltoides*, chromatic contrast (as perceived by bees) in *L. ircutianum*, and chroma in *V. chamaedrys* (Detailed information in Table S2). Thus, for specific traits across certain species, soil moisture conditions, by governing mass flow rates and root nutrient assimilation (Plett *et al*. 2020), dictate whether additional nutrients yield either positive or negative phenotypic outcomes.

While *A. arenosa* experienced a reduction in chroma under nutrient enrichment, ray florets of *L. ircutianum* surprisingly experienced an increase in chroma and chromatic contrast (as perceived by bees and flies). This increase was also mirrored under water scarcity, which similarly elevated chroma and chromatic contrast (in flies in *L. ircutianum*). The contrasting responses of these two species to nutrient enrichment likely stem from fundamental differences in their floral pigmentation pathways. Unlike polymorphic species like *A. arenosa*, *L. ircutianum* features only white ray florets (and yellow disc florets) that typically lack visible anthocyanins due to mutations or down-regulation of genes within the flavonoid pathway (Ahmad *et al*. 2023). However, severe abiotic stress can disrupt this status quo. For instance, drought has been shown to induce pigmentation in otherwise white-flowered lineages by reactivating latent anthocyanin production pathways (Vaidya *et al*. 2018). It is highly possible that both nutrient enrichment and water scarcity acted as independent environmental stressors in *L. ircutianum*, triggering the activation of these dormant biosynthetic pathways and resulting in the observed increases in floral chroma.

### 4.2. Mediators of effects of Abiotic factors on Pollinator Visitation

While investigating how abiotic factors alter floral traits provides insights into individual species response to global change, evaluating how these shifts scale up to influence pollinator attraction is crucial for understanding broader plant community dynamics. In our study, the effects of abiotic factors on pollinator visitation were evident in only three of the five focal species. While a large proportion of these effects could be explained by the traits measured in our study, significant direct effects of abiotic factors remain unexplained, both across species and within individual species. These effects can likely be attributed to the unmeasured floral rewards and volatiles which are well-known to impact pollinator behavior (e.g., Muhlemann, Klempien, and Dudareva 2014, Vaudo *et al*. 2022), but were not studied by us due to technical constrains.

A second potential limitation involves floral phenology. Both nutrient enrichment and water scarcity are known to alter flowering phenology. Nitrogen addition has been shown to delay flower in certain functional groups while advancing it in others (Zhang *et al*. 2014) while water scarcity has been found to significantly alter both flowering phenology and floral lifespan (reviewed in Brunet *et al*. 2025). Although we ensured that pollinator observations were conducted when all treatments reached peak flowering, treatment-specific variations in phenological timing may have meant that plants in certain treatments were at slightly more advanced flowering stages than others. Furthermore, we cannot verify whether our observational windows during peak plant flowering synchronized with peak activity periods of local wild pollinators (which were dominated by hoverflies and solitary bees). While using reared pollinator colonies would eliminate this temporal mismatch, doing so would less representative of natural pollination dynamics.

In *D. deltoides* and *P. argentea*, we did not detect any significant link connecting abiotic factors to flower visitation. This is in line with broad scale meta-analysis demonstrating that environmental stressors like water scarcity do not always result in predictable decline in pollinator visitation rates (Kuppler and Kotowska 2021). Despite broad effects of abiotic factors on plant traits, none of the measured traits acted as effective intermediaries for visitation in these species nor were there any significant links between abiotic factors and visitation indicating effects of some unmeasured traits. This lack of response cannot be attributed to differences in their dominant pollinators or pollinator visitation rates as they were visited by similar pollinators in similar frequencies as the other 3 species. Therefore, our findings indicate that pollinators could be tolerant of changes in floral trait expression in some species (in our case, *D. deltoides* and *P. argentea*) but not in other species.

Across the three species where effect of abiotic factors on pollinator visitation was evident, nutrient enrichment influenced pollinator visitation by altering plant height, flower number, floral area, chromatic contrast (in flies) and chroma. This strong trait-mediated relationship is at odds with several previous studies suggesting that nutrient-driven changes in plant morphology have no impact on visitation, which is instead primarily dictated by floral rewards such as nectar volume, pollen quantity, and amino acid concentrations (Vaudo *et al*. 2022, Leach *et al*. 2023). Our findings reaffirm that changes in floral visual traits do influence pollinator visitation, and they can be better understood by considering how pollinators forage. Pollinators frequently rely on external morphology as reliable visual proxies for these internal rewards (Tavares, Freitas, and Gaglianone 2016). For example, bees utilize overall flower size at longer distances and chromatic contrast at shorter ranges to evaluate potential floral rewards before landing (Ortiz *et al*. 2021). Given the positive and negative responses of these visual traits to nutrient enrichment, alterations in pollinator attraction could serve as a major driver of shifting plant community composition. This aligns with recent experimental evidence demonstrating that nutrient enrichment alters grassland community composition by shifting the abundance of flowers displaying different visual colours (Nelson, Borer, and Seabloom 2025).

Water scarcity also exerted effects on pollinator visitation, operating through plant height, floral area, chromatic contrast (as perceived by flies) and chroma. While experimental studies confirm that drought-stressed plants with reduced floral displays receive fewer pollinator visits (e.g., Rering *et al*. 2020, Rose-Person, Santiago, and Rafferty 2025), large-scale meta-analysis revealed significant heterogeneity in these outcomes (Kuppler and Kotowska 2021). It should be noted that this heterogeneity can be resolved in future studies through certain improvements in experimental methodology (Brunet *et al*. 2025) such as exposing plants of different experimental treatments separately to pollinators which would be more representative of natural conditions. In our study, the direct negative effect of water scarcity on pollinator visitations in *L. ircutianum* and *V. chamaedrys* is likely explained by unmeasured reductions in floral rewards as drought consistently reduces nectar volume (Kuppler and Kotowska 2021) which strongly deters pollinator visitation.

Intriguingly, *L. ircutianum* plants increased chroma under both nutrient enrichment and water scarcity, despite this elevated color intensity having a negative effect on pollinator visitation. This finding can be understood through the lens of *correlated selection*. While pollinators impose strong selection on floral color (Hopkins and Rausher 2012), color traits do not evolve in a vacuum as they also represent vital physiological adaptations to abiotic stressors like drought, UV radiation and land-use (Binkenstein, Renoult, and Schaefer 2013, Dalrymple *et al*. 2020). This is because the pigments responsible for floral color such as anthocyanins play dual roles in UV absorption, scavenging Reactive Oxygen Species (ROS) and maintaining osmotic balance (reviewed in Dabravolski and Isayenkov, 2023). Therefore, the physiological demand of stress tolerance may ultimately override or constrain selection for pollinator preference.

In *A. arenosa* and *L. ircutianum*, the combination of nutrient enrichment and water scarcity also influenced pollinator visitation, particularly by variations in plant height, floral area, chromatic contrast (as perceived by flies) and chroma. The combination of both stressors elicited both positive and negative responses depending entirely on species and trait. This complexity highlights an important gap in the limited existing literature in population ecology focusing on multi-stressor experiments. For instance, Akter and Klecka (2022) set up an experiment exploring how nutrient and water availability and their combination influence flower visitation. Their statistical analyses however solely tested the independent main effects of individual factors on pollinator visitation, and not the interaction, despite testing the interaction effects on plant traits. They reported that while water availability altered visitation, nitrogen supply had no discernible effect. By explicitly modeling the interaction term for pollinator visitation as well, our study demonstrates that water scarcity does not simply operate independently and interacts with nutrient enrichment in unique ways to affect pollinator visitation.

## 5. Acknowledgements

This study was supported by the Grantová Agentura České Republiky (project number 22- 00761S), partially funded by the long-term research development project (no. RVO 67985939) of the Akademie Věd České Republiky, and by Institutional support for science and research from the Ministry of Education, Youth, and Sports of the Czech Republic. We thank David Cmíral and Miroslav Chmelař for their assistance in both the preparation and maintenance of the experimental setup. We thank Dana Parysova for her technical assistance with the project. We thank our Open Science research interns of 2024, Filip Kreuter and Tereza Kratochvílová for their assistance in processing of plant tissue samples. We thank the members of the POPEKOL group for their insightful comments on the development of our manuscript. We also wish to disclose that AI-based tools were utilized to aid in literature gathering, assist with statistical analysis, and refine the tone of certain sections; the authors remain fully responsible for the content and interpretation.

## 7. Author contributions

MM, DT and ZM planned and designed the research study and experiment. MM prepared the multi-species experimental setup following which along with BP, performed the sampling and processing of floral traits. BP conducted the pollinator visitation experiment for the individual species. MM analyzed the corresponding data and performed all statistical analyses with input from DT and ZM. MM drafted the manuscript, with substantial suggestions, inputs and revisions from DT and ZM.

## 8. Data availability

The data that supports the findings of this study will be made openly available in a Zenodo repository upon acceptance along with the relevant software scripts being attached in GitHub. Prior to acceptance, data will be made available to reviewers on request to the corresponding author.

