## Supporting Information for "Interactive effects of Nutrient Enrichment and Water Scarcity on Floral signaling and visitation"

The following Supporting Information is available for this article:

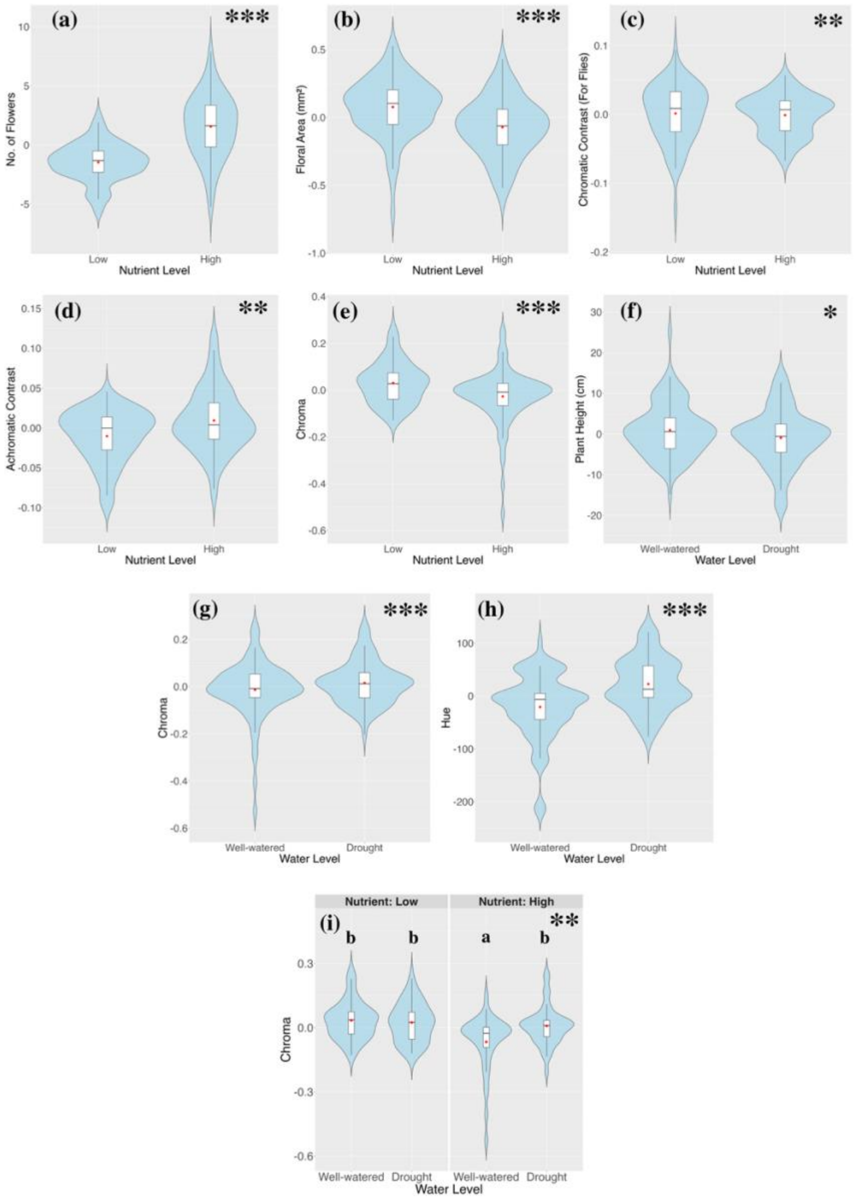


**Fig. S1.** Violin plots illustrating the individual and interactive effects of nutrient enrichment and water limitation on floral traits of in all species. Only significant plots are shown; the significance level of the effect of the plotted factor is indicated in the top right corner (*$P$= 0.05; **$P$≤ 0.01; ***$P$ ≤ 0.001). The functional trait studied is denoted on the *Y*-axis while factor is denoted in the *X*-axis. Trait values are presented as residuals derived from linear models accounting for species identity (Trait ~ Species). Different lowercase letters above the violins indicate statistically significant differences between treatment combinations based on Tukey's HSD post-hoc test ($P$≤ 0.05).

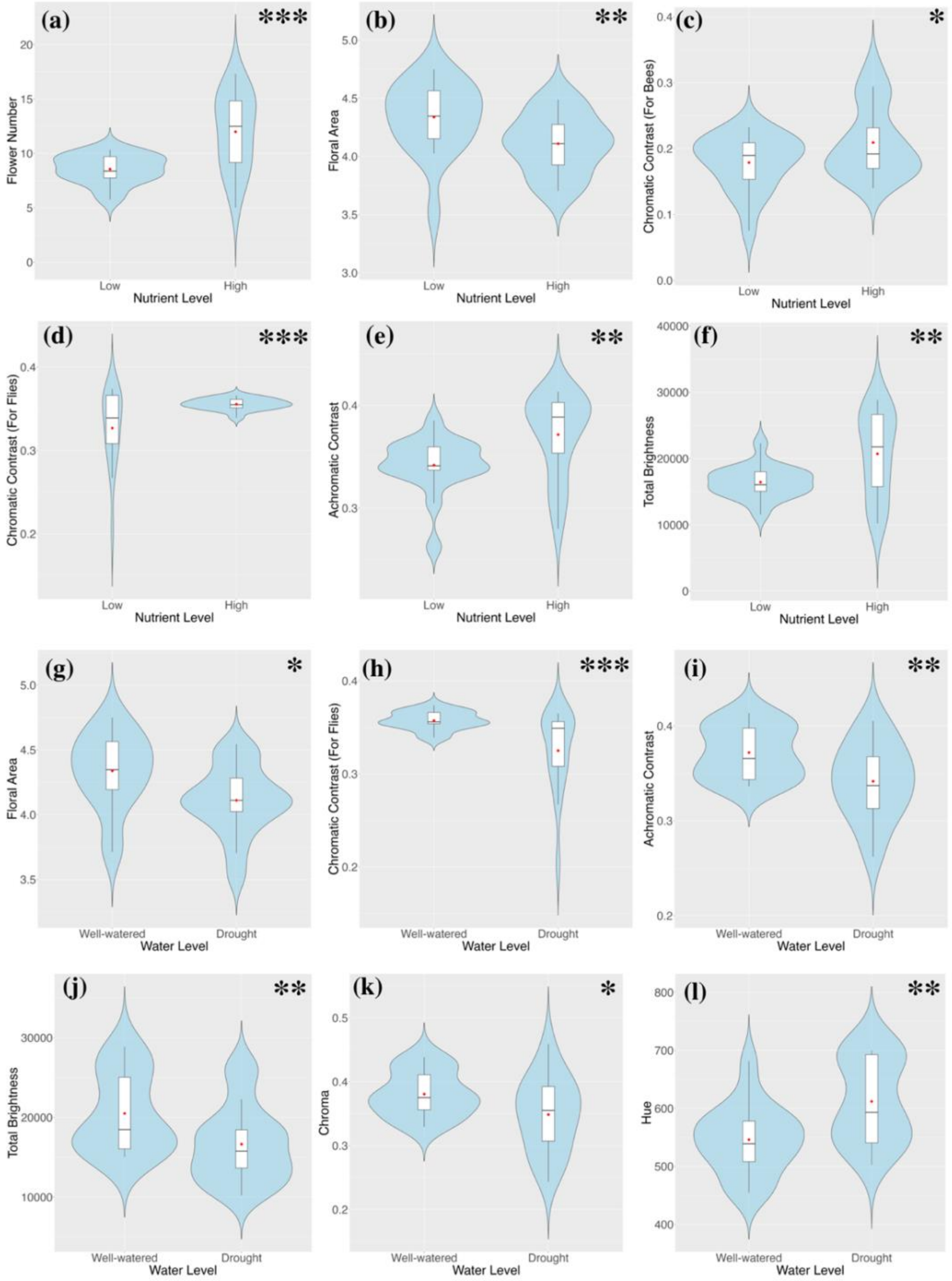


**Fig. S2.** Violin plots illustrating the individual effects of nutrient enrichment and water limitation on floral traits of *A. arenosa*. Only significant plots are shown; the significance level of the effect of the plotted factor is indicated in the top right corner (*$P$= 0.05;**$P \leq$ 0.01;***$P \leq$ 0.001). The functional trait studied is denoted on the *Y*-axis while factor is denoted in the *X*-axis.

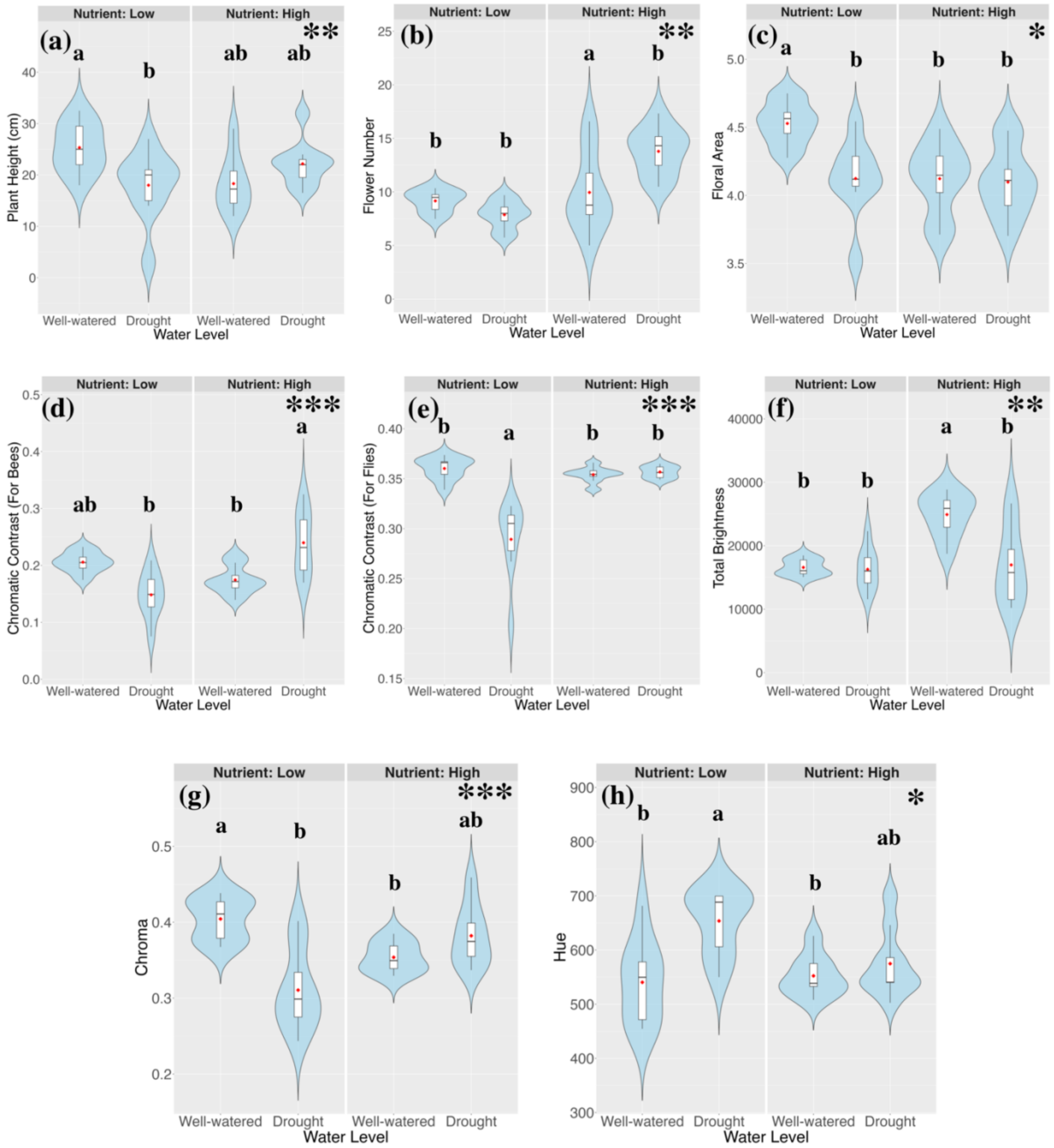


**Fig. S3.** Violin plots illustrating the interactive effects of nutrient enrichment and water limitation on floral traits of *A. arenosa*. Only significant plots are shown; the significance level of the effect of the plotted interaction is indicated in the top right corner (*$P$= 0.05;**$P \leq$ 0.01;***$P \leq$ 0.001). The functional trait studied is denoted on the *Y*-axis while factor is denoted in the *X*-axis. Different lowercase letters above the violins indicate statistically significant differences between treatment combinations based on Tukey's HSD post-hoc test ($P \leq$ 0.05).

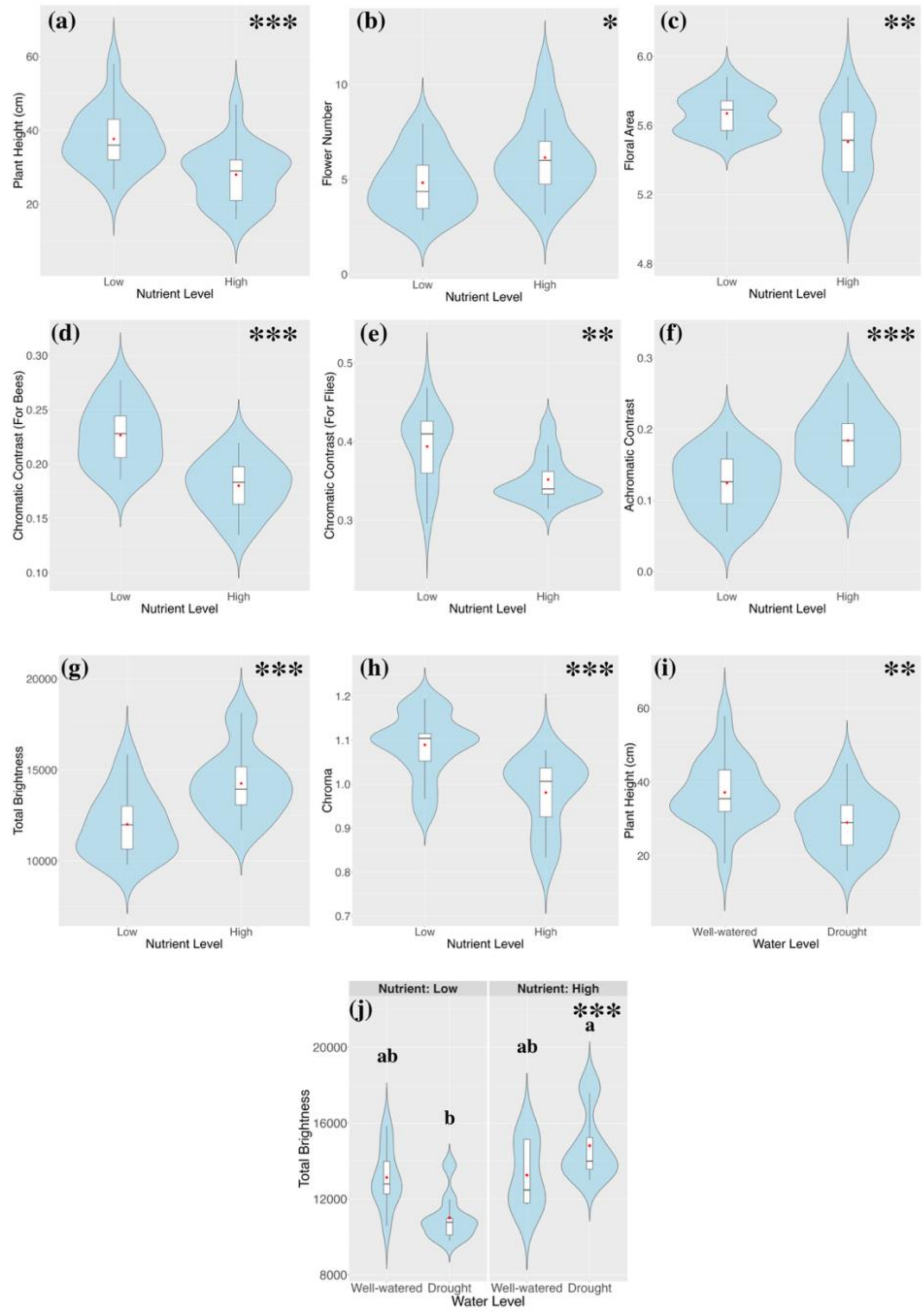


**Fig. S4.** Violin plots illustrating the individual and interactive effects of nutrient enrichment and water limitation on floral traits of *D. deltoides*. Only significant plots are shown; the significance level of the effect of the plotted factor or their interaction is indicated in the top right corner (*$P$= 0.05;**$P \leq$ 0.01;***$P \leq$ 0.001). The functional trait studied is denoted on the *Y*-axis while factor is denoted in the *X*-axis. Different lowercase letters above the violins indicate statistically significant differences between treatment combinations based on Tukey's HSD post-hoc test ($P \leq$ 0.05).

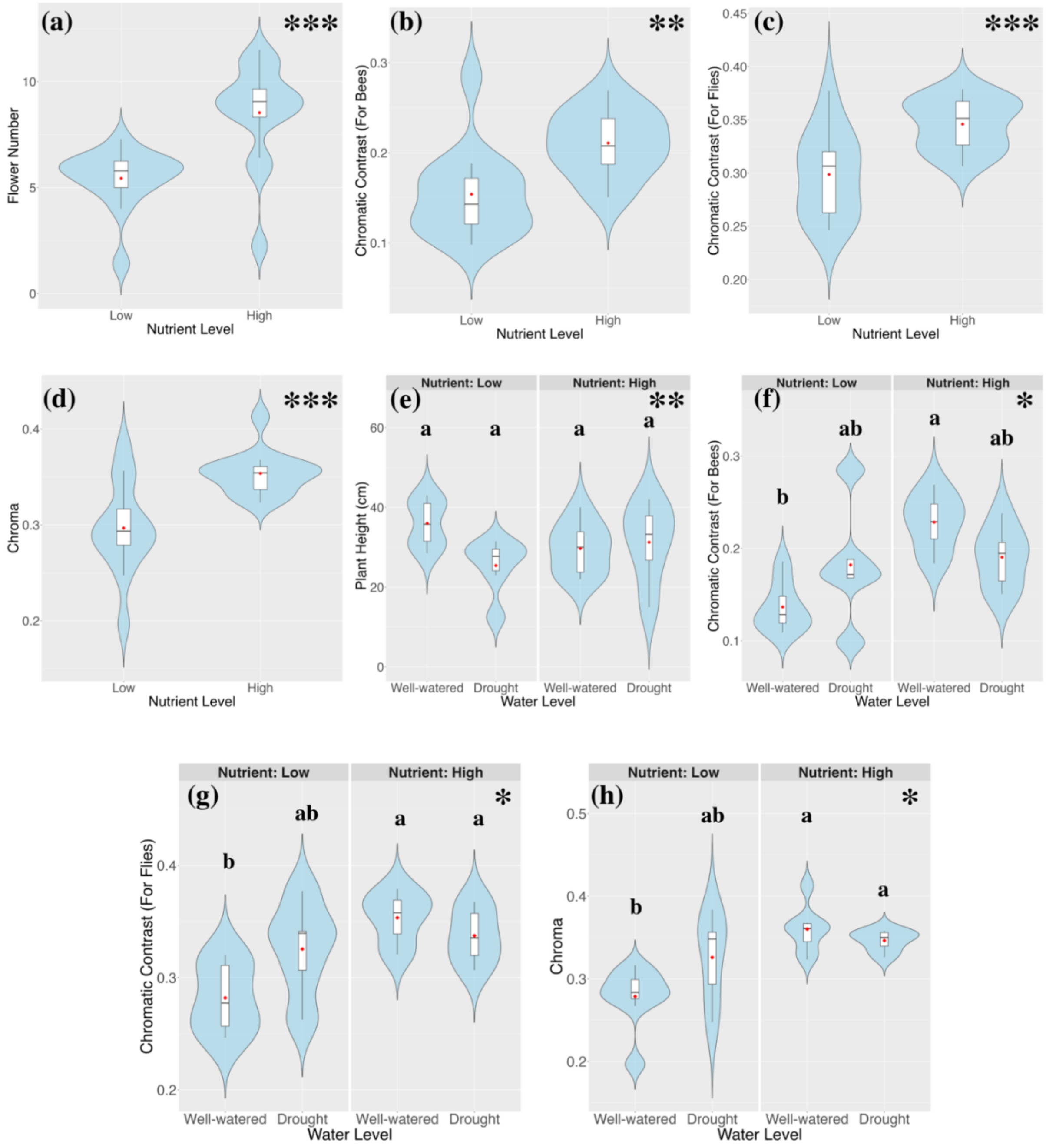


**Fig. S5.** Violin plots illustrating the individual and interactive effects of nutrient enrichment and water limitation on floral traits of *L. ircutianum*. Only significant plots are shown; the significance level of the effect of the plotted factor or their interaction is indicated in the top right corner (**P*= 0.05;***P*≤ 0.01;****P* ≤ 0.001). The functional trait studied is denoted on the *Y*-axis while factor is denoted in the *X*-axis. Different lowercase letters above the violins indicate statistically significant differences between treatment combinations based on Tukey's HSD post-hoc test (*P*≤ 0.05).

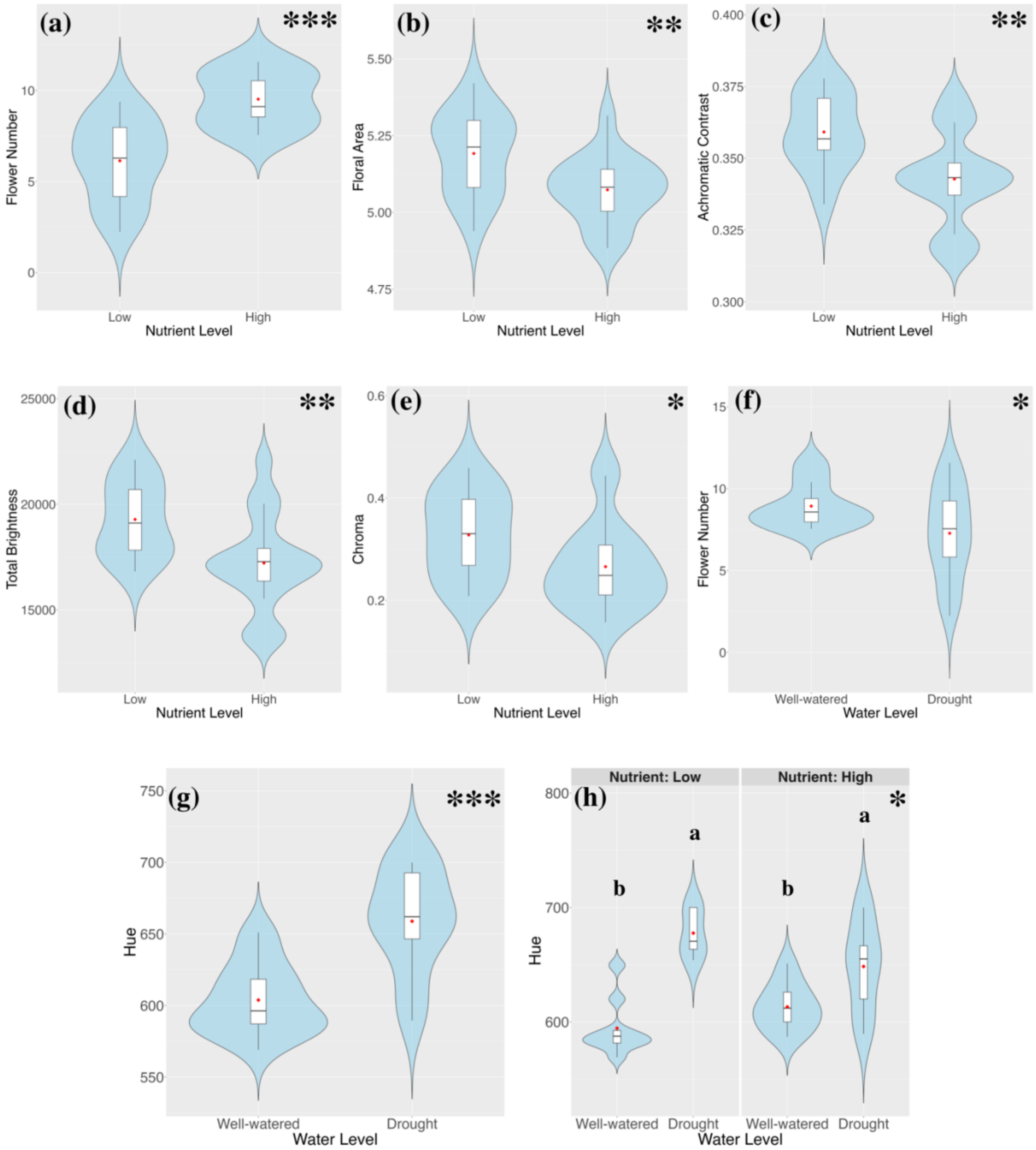


**Fig. S6.** Violin plots illustrating the individual and interactive effects of nutrient enrichment and water limitation on floral traits of *P. argentea*. Only significant plots are shown; the significance level of the effect of the plotted factor or their interaction is indicated in the top right corner ( $*P=0.05$; $**P\leq 0.01$; $***P \leq 0.001$). The functional trait studied is denoted on the *Y*-axis while factor is denoted in the *X*-axis. Different lowercase letters above the violins indicate statistically significant differences between treatment combinations based on Tukey's HSD post-hoc test ($P\leq 0.05$).

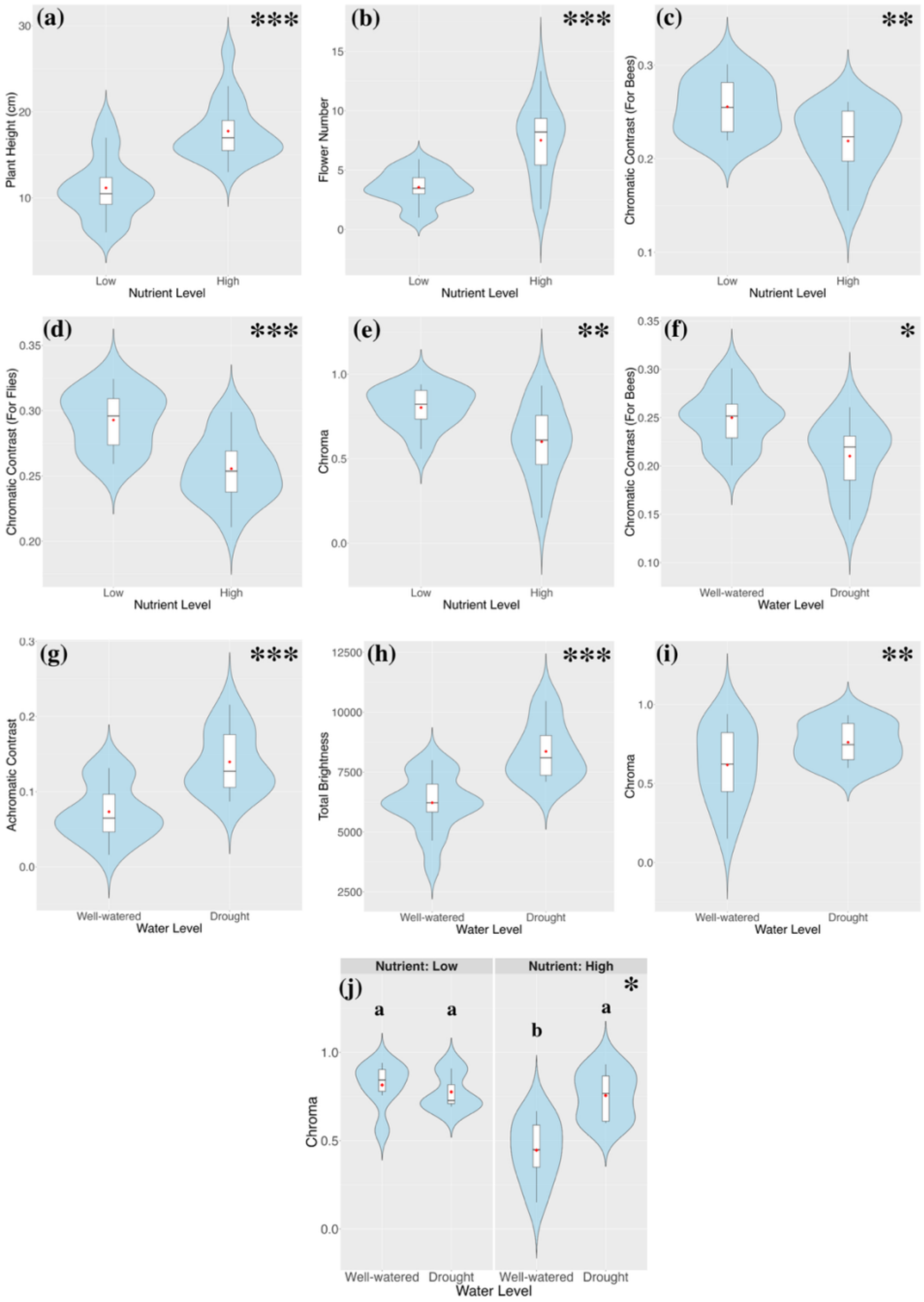


**Fig. S7.** Violin plots illustrating the individual and interactive effects of nutrient enrichment and water limitation on floral traits of *V. chamaedrys*. Only significant plots are shown; the significance level of the effect of the plotted factor or their interaction is indicated in the top right corner (*$P$= 0.05;**$P \leq 0.01$;***$P \leq 0.001$). The functional trait studied is denoted on the *Y*-axis while factor is denoted in the *X*-axis. Different lowercase letters above the violins indicate statistically significant differences between treatment combinations based on Tukey's HSD post-hoc test ($P \leq 0.05$).

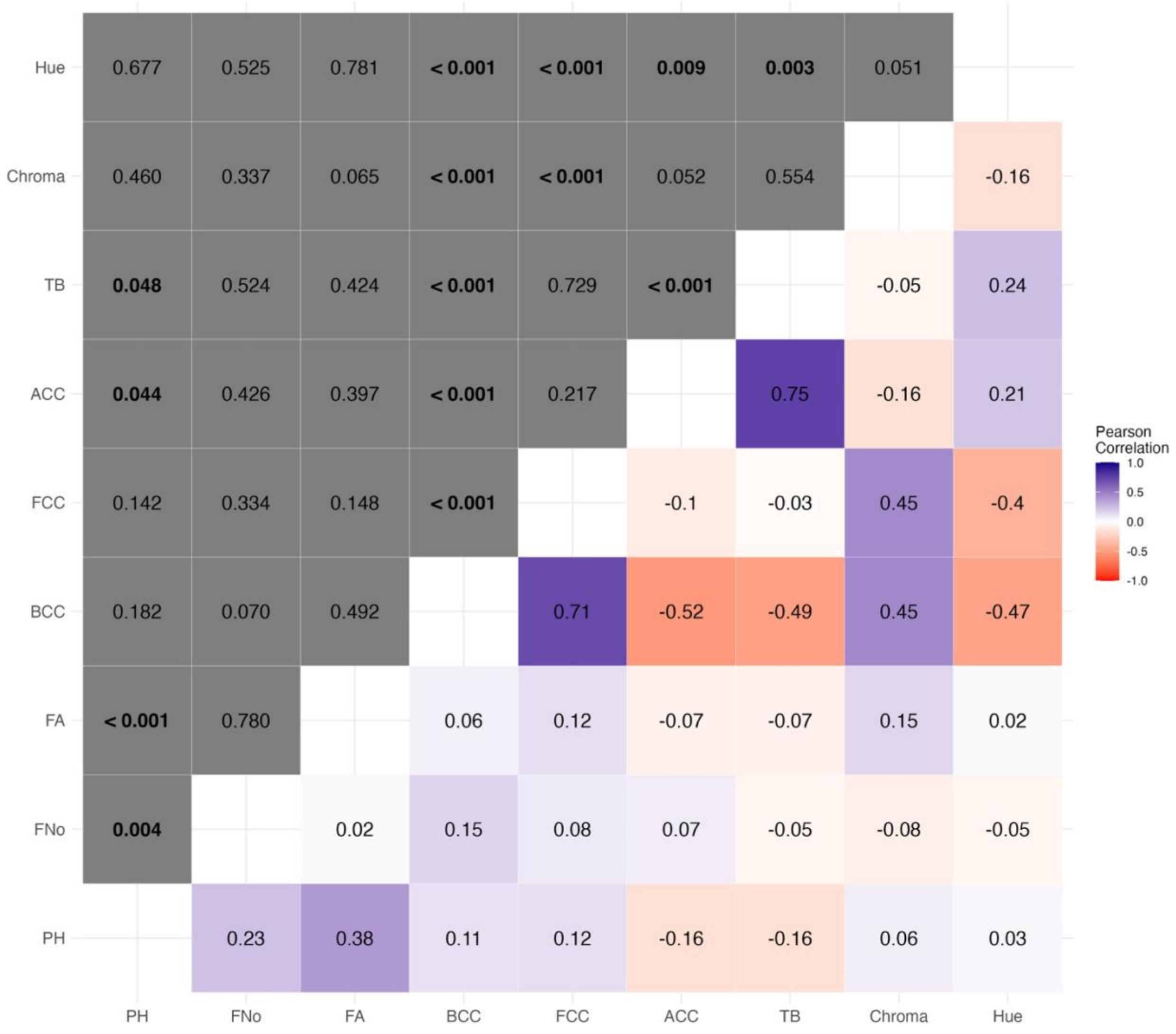


**Fig. S8.** Correlation heatmap of studied plant traits with the coefficient (r-value) of the Pearson correlation test of a trait pair indicated in the lower right triangle and its p-value indicated in the upper left triangle. PH, Plant Height; FNo, Flower Number; FA, Floral Area; BCC, Chromatic Contrast for Bees; FCC, Chromatic Contrast for Flies; TB, Total Brightness.

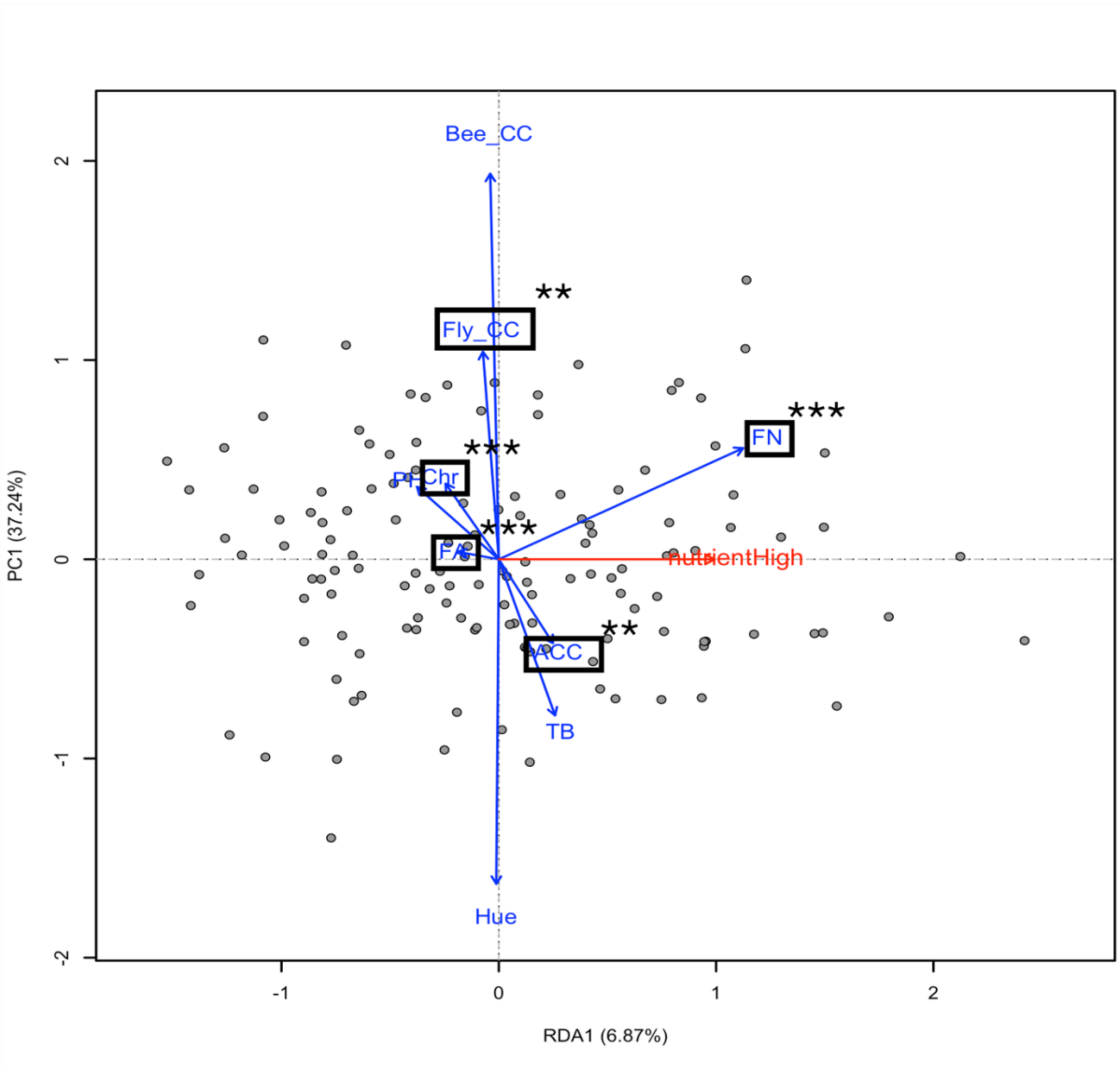


**Fig. S9.** RDA plot representing effects of nutrient enrichment on studied plant traits. Significance level of the effect of nutrient enrichment on trait has been indicated in the top right corner of the trait (**$P \leq 0.01$; ***$P \leq 0.001$.). PH, Plant Height; FN, Flower Number; FA, Floral Area; Bee_CC, Chromatic Contrast for Bees; Fly_CC, Chromatic Contrast for Flies; TB, Total Brightness; Chr, Chroma.

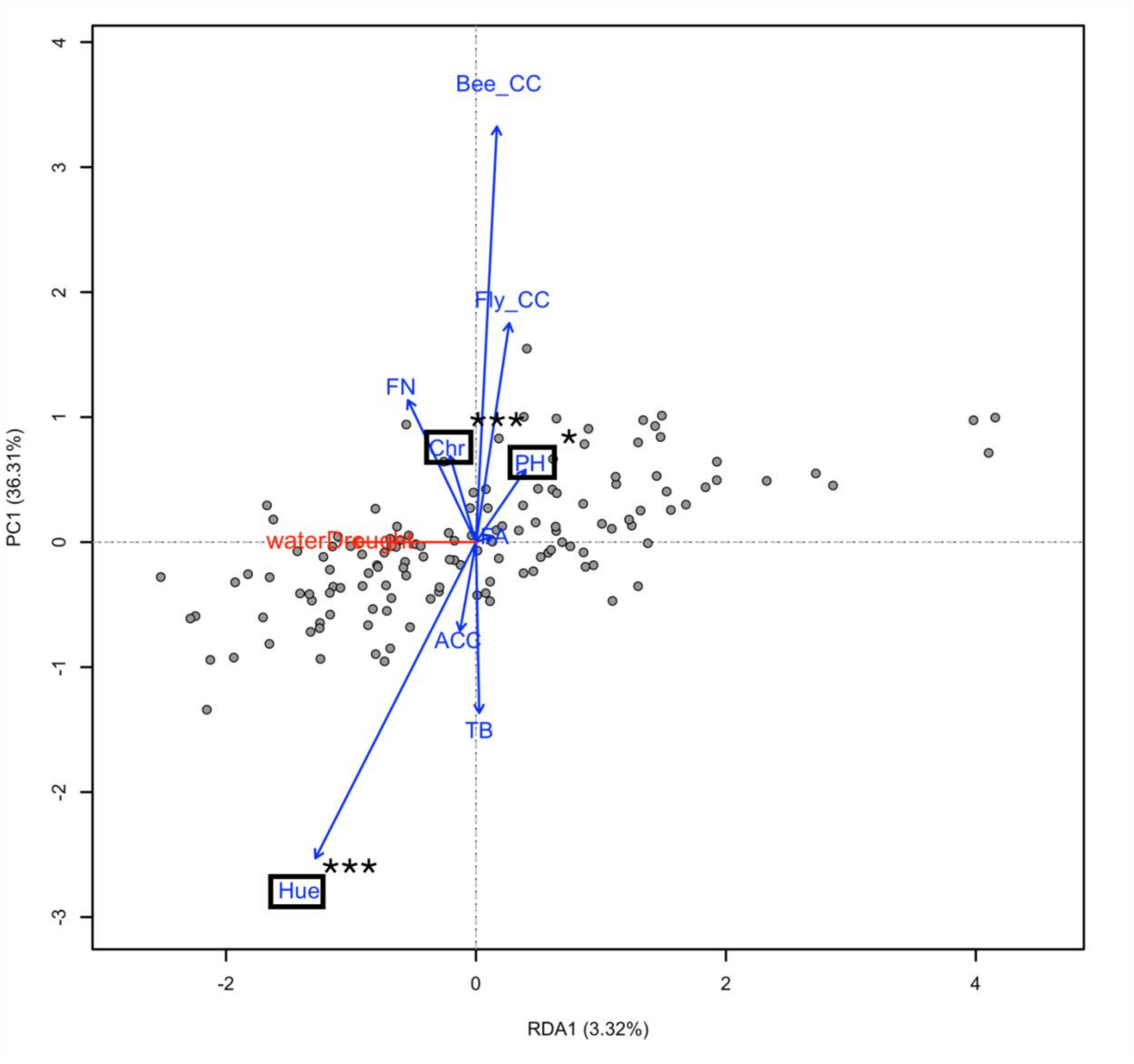


**Fig. S10.** RDA plot representing effects of water scarcity on studied plant traits. Significance level of the effect of water scarcity on trait has been indicated in the top right corner of the trait (*$P$= 0.05;**$P$≤ 0.01;***$P$ ≤ 0.001.). PH, Plant Height; FN, Flower Number; FA, Floral Area; Bee_CC, Chromatic Contrast for Bees; Fly_CC, Chromatic Contrast for Flies; TB, Total Brightness; Chr, Chroma.

| Species | Fisher's C | df | p-value |
|---|---|---|---|
| *A. arenosa* | 40.172 | 44 | 0.636 |
| *D. deltoides* | 42.312 | 64 | 0.983 |
| *L. ircutianum* | 21.965 | 32 | 0.908 |
| *P. argentea* | 36.484 | 58 | 0.988 |
| *V. chamaedrys* | 48.435 | 60 | 0.858 |

**Table. S1.** Goodness-of-fit statistics for species-specific piecewise Structural Equation Models.

| Species | Trait | Effect of interaction |
|---|---|---|
| *A. arenosa* | Plant Height | Negative effect of water scarcity only pronounced in low-nutrient conditions |
| *A. arenosa* | Flower Number | Nutrient enrichment leads to increase in flower number regardless of moisture conditions with plants experiencing both nutrient enrichment and water scarcity having the highest flower number. Positive effects of water scarcity more pronounced under nutrient enrichment. |
| *A. arenosa* | Floral area | Nutrient enrichment leads to decrease in floral area regardless of moisture conditions. Negative effects of water scarcity more pronounced in low nutrient condition. |
| *A. arenosa* | Chromatic Contrast (as perceived by bees) | Water scarcity leads to decrease in chromatic contrast in low nutrient condition but increased chromatic contrast under nutrient enrichment. |
| *A. arenosa* | Chromatic Contrast (as perceived by flies) | Water scarcity leads to decrease in chromatic contrast in low nutrient condition but not under nutrient enrichment. |

| | | |
|---|---|---|
| *A. arenosa* | Total Brightness | Nutrient enrichment leads to increase in total brightness but only under well-watered conditions. |
| *A. arenosa* | Chroma | While chroma is highest in low-nutrient, well-watered conditions, negative effect of water scarcity is present only in low-nutrient conditions. |
| *A. arenosa* | Hue | Water scarcity leads to increase in hue but only under low nutrient conditions. |
| *D. deltoides* | Total Brightness | Nutrient enrichment leads to increase in total brightness but only under water scarcity |
| *L. ircutianum* | Chromatic Contrast (as perceived by bees) | Nutrient enrichment leads to increase in chromatic contrast but only under well-watered conditions. |
| *L. ircutianum* | Chromatic Contrast (as perceived by flies) | Nutrient enrichment leads to increase in chromatic contrast regardless of moisture conditions. |
| *L. ircutianum* | Chroma | Nutrient enrichment leads to increase in chroma regardless of moisture conditions. |
| *P. argentea* | Hue | Water scarcity leads to increase in hue regardless of nutrient conditions. |
| *V. chamaedrys* | Chroma | Nutrient enrichment leads to decrease in chroma but only in well-watered conditions. |

**Table. S2.** Detailed results for effects of interaction between nutrient enrichment and water scarcity in individual species.

| Species | Family | Ellenberg - Moisture | Ellenberg - Nutrient |
|---|---|---|---|
| *A. arenosa* | Brassicaceae | 4 | 3 |
| *D. deltoides* | Caryophyllaceae | 4 | 3 |
| *L. ircutianum* | Asteraceae | 5 | 5 |
| *P. argentea* | Rosaceae | 2 | 3 |
| *V. chamaedrys* | Plantaginaceae | 5 | 6 |

**Table. S3.** Ellenberg indicator moisture and nutrient values for individual species
